# Divine: Prioritizing Genes for Rare Mendelian Disease in Whole Exome Sequencing Data

**DOI:** 10.1101/396655

**Authors:** Changjin Hong, Jean R. Clemenceau, Yunku Yeu, TaeHyun Hwang

## Abstract

**Motivation:** Recent studies showed that a phenotype-driven analysis of whole exome sequencing (WES) could provide more accurate and clinically relevant genetic variants.

**Results:** We develop a computational tool called Divine that integrates patients’ phenotype(s) and WES data with 30 prior biological knowledge (e.g., human phenotype ontology, gene ontology, pathway database, protein-protein interaction networks, pathogenicity by the amino acid change due to polymorphism, and hot-spot protein domains) to prioritize potential disease-causing genes. In a retrospective study with 22 real and four simulated data set, Divine ranks the same pathogenic genes confirmed by the original studies 5th on average out of a thousand of mutated genes and outperforms existing state-of-the-art methods.

**Availability:** https://github.com/hwanglab/divine

**Supplementary information:** Supplementary Document is attached at the end of the page.

## 1 Introduction

Recently, whole-exome sequencing (WES) has become a viable option for Mendelian diseases diagnosis. As more genes are considered for testing, it becomes substantially challenging to interpret and identify mutations responsible for patients’ illnesses among thousands of variants in the massively high-throughput data.

It has been shown that prioritizing the gene known to be associated with a patient’s phenotype [Köhler 2017] facilitates variant interpretation. We can focus on a gene set by matching the patient’s symptomatology to the list of known disorders and estimate the significance of each disease match [Sifrm 2013, Javed 2014, Zemojtel 2014]. Furthermore, the disease-gene association can be indirectly expanded to the other species [Robinson 2014]. The phenotype information incorporates genetic variants in either a Bayesian framework [Javed 2014] or Random Forest [Antanaviciute 2015] voting model. However, the existing methods are limited to only the prior knowledge domain still far from completeness, and we are also cautious not to over-amplify it due to redundant heterogeneous databases. It is hard to find a freely available standalone package that provides both all annotations necessary to comply with ACMG guidelines and prioritized genes for a rapid diagnosis.

We develop a standalone software, *Divine*, to prioritize the underlying mutated genes of rare hereditary disorders. Divine accepts either Human Phenotype Ontology (HPO) IDs manifesting patient clinical features or any VCF file generated from WES reads. Divine employs semantic similarity between patient phenotype and known diseases registered in the most reliable and up-to-date database. It also incorporates both Gene ontology and Kyoto Encyclopedia of Genes and Genomes (KEGG) pathway enrichment to discover genes previously never reported to be associated with any disease. To improve specificity, we introduce two new variant annotations: 1) pathogenicity of each protein domain and 2) pathogenic likelihood ratio due to nonsynonymous SNV trained from variant datasets like ClinVar, HGMD [Stenson 2017], and 1000 genome project.

## 2 Methods

The workflow starts by computing the semantic similarity of the patient’s phenotypic terms to each known disease catalog with the help of HPO summarizing OMIM and Orphanet, covering a total of 5,528 diseases, 4,845 genes, and 6,090 HPO terms. Then, a set of term-to-term similarity scores of the query HPOs for each disease is derived from maximal term-to-term scores [Schlicker 2008] with a modification to alleviate the issue where symptoms related to disease are incomplete or sparse. The phenotypic matching score (*P*_i_) is given to each gene associated with the disease.

Given a patient VCF file, we predict the genetic damage of each gene harboring variants based on multiple orthogonal annotations. Divine establishes an open source variant annotation tool, *Varant* (See S. Table 1). We substantially improve the original work by adding eight new annotations. As a result, 30 total annotations are available.

Divine assigns a variant pathogenic score (*G*_i_) considering the following criteria: 1) the variant MAF, 2) zygosity per transcript (e.g., heterozygous, homozygous, or homo compound heterozygous), 3) a functional impact of the variant within the transcript (e.g., null variant causing a loss of function and splice site/enhancer/silencer), 4) whether a variant locates within a mutational hotspot or critical and well-established functional domain, and 5) a pathogenic likelihood ratio with respect to an amino acid change inferred from two in silico prediction score distributions built from known benign variants and pathogenic variants, respectively. See Section 4.2 in Supp. Document.

**Fig 1.**
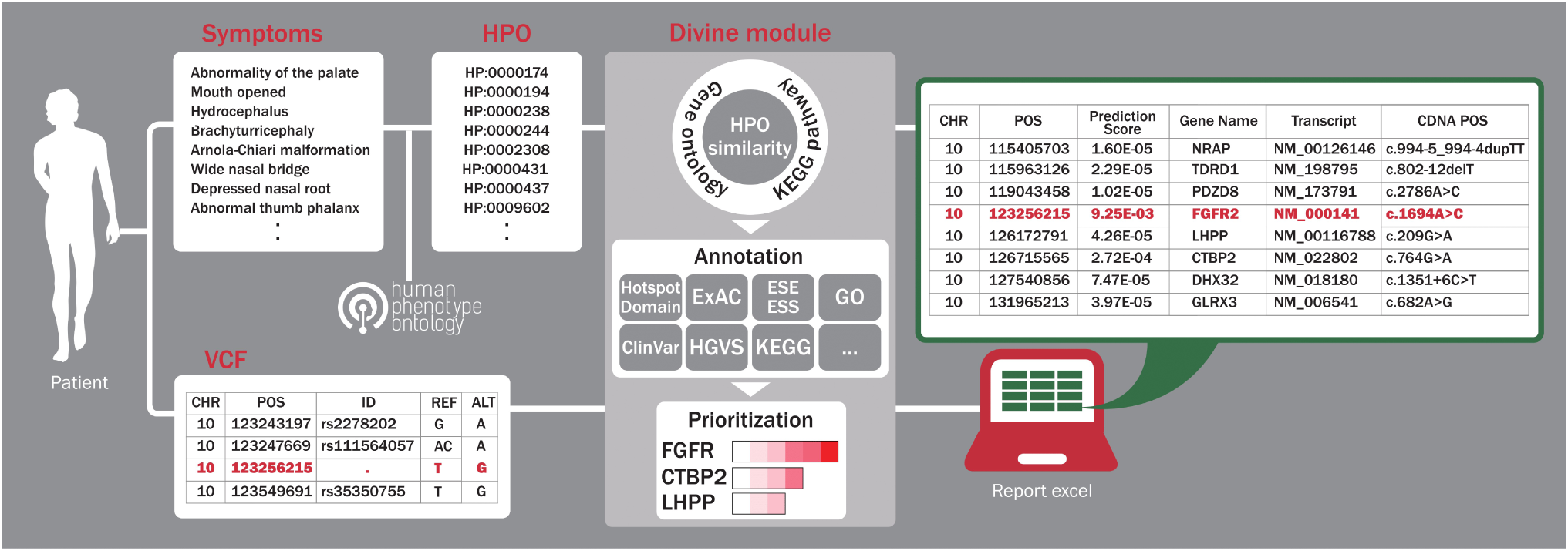
*Divine (https://github.com/hwanglab/divine) takes a VCF or/and HPO IDs. Symptom to ID conversion can be done at https://hpo.jax.org/app or https://mseqdr.org/search_phenotype.php. An output table contains 30 variant/gene-level annotations with prioritized gene ranking by pathogenicity (prediction score).*

Both phenotypic score (*P*_i_) and variant score (*G*_i_) described above are combined into a single value (*y*_i_) [Javed 2014]. Genes with higher *P*_i_ indirectly populate a new *P*_i_ to the genes with higher *G*_i_, but its *P*_i_ is not available through either KEGG pathway memberships or Gene ontology similarity. Finally, the *y*_i_ is propagated in a gene interaction network connecting 19,035 total genes using a heat diffusion kernel [Yang 2007] (See Section 4.5 in Supp. Document).

## 3 Results

Divine is compared with five other methods, *Phen-Gen_v1, Exomiser_v7.2*, *Exomiser_v10*, *eXtasy*, and *PhenIx*, and is tested with three simulated samples and 23 real patient VCF files [Antanaviciute 2015] studied between 2012 and 2016, covering a broad spectrum of rare Mendelian diseases (average 10.8 HPO terms).

Both Divine (AUC score: 0.959) and *Exomiser_v10* (0.954) outperform the other methods, *Phen-Gen_v1 (0.482)*, *Exomiser_v7.2* (0.577), *eXtasy* (0.5), and *PhenIX* (0.728) (Refer to S. Table 3 and S. Fig 3). Divine is more specific than *Exomiser_v10*, and it reports the genes of interest within 21st from the top for 96.2% of the samples vs. 88.5% for *Exomiser_v10*. In an experiment to test a new disease-to-gene discovery (Section 5.4 and S. Table 5 in Suppl. Document), Divine ranks all 4 genes harboring pathogenic variants within 6^th^ from the top via either Gene ontology, KEGG pathway, or PPI network where each gene enriched by its partner is associated with a disease with similar phenotypes. Finally, in a robustness test where we randomly introduce irrelevant extra HPO terms, Divine prioritizes the disease-causing genes on average rank 8^th^ from the top for 22 out of 26 cases (84.6%).

## 4 Conclusion

Divine can facilitate molecular diagnosis with clinical whole exome sequencing data swamped by a significant number of variants. It harmonizes the most comprehensive and extensible annotations to find genes harboring disease-causing variants. Divine supports a discovery mode via phenotype enrichment to search for new disease-associated genes.

